# Megakaryocyte maturation involves activation of the adaptive unfolded protein response

**DOI:** 10.1101/2023.07.26.550730

**Authors:** M. Faiz, M.L Kalev-Zylinska, C. Dunstan-Harrison, D.C Singleton, M.P. Hay, E.C. Ledgerwood

## Abstract

Endoplasmic reticulum stress triggers the unfolded protein response (UPR) to promote cell survival or apoptosis. Transient endoplasmic reticulum stress activation has been reported to trigger megakaryocyte production, and UPR activation has been reported as a feature of megakaryocytic cancers. However, the role of UPR signaling in megakaryocyte biology is not fully understood. We studied the involvement of UPR in human megakaryocytic differentiation using PMA (phorbol 12-myristate 13-acetate)-induced maturation of megakaryoblastic cell lines and thrombopoietin-induced differentiation of human peripheral blood-derived progenitors. Our results demonstrate that an adaptive UPR is a feature of megakaryocytic differentiation, and that this response is not associated with ER stress-induced apoptosis. Differentiation did not alter the response to the canonical endoplasmic reticulum stressors DTT or thapsigargin. However, thapsigargin, but not DTT, inhibited differentiation, consistent with the involvement of Ca^2+^ signaling in megakaryocyte differentiation.

## Introduction

Megakaryocytes are bone marrow cells that produce platelets. The differentiation of hematopoietic stem cells (HSC) to megakaryocytes (megakaryopoiesis) and subsequent platelet release (thrombopoiesis) is controlled by a number of transcription factors and cytokines. HSC undergo multiple rounds of endomitosis to produce large polyploid mature megakaryocytes with an extensive internal membrane reservoir for platelet formation. Recent reports suggest that activation of the endoplasmic reticulum (ER) stress response is a feature of both normal megakaryopoiesis and thrombopoiesis (Kovuru et al., 2020; Lopez et al., 2013; Morishima & Nakanishi, 2016), and abnormal megakaryopoiesis in *CALR*-mutated myeloproliferative neoplasms (El-Khoury et al., 2020; Fosselteder et al., 2023; Jutzi et al., 2023; Nam et al., 2019; Prins et al., 2020).

The ER is involved in folding and post-translational modification of secreted and transmembrane proteins, storage of calcium, and lipid membrane biogenesis, with disruptions of these processes leading to ER stress (Braakman & Hebert, 2013; Clapham, 2007; Fagone & Jackowski, 2009). In response to ER stress, eukaryotic cells trigger the unfolded protein response (UPR) that initially helps cells to overcome the stress by upregulating the expression of chaperones to promote folding, repressing mRNA translation to temporarily inhibit global protein synthesis, and initiating ER-associated degradation (ERAD) pathways (adaptive UPR) (Feral et al., 2021; Hetz et al., 2020). If the adaptive UPR is unable to restore protein homeostasis, UPR signaling events may cause apoptosis (terminal UPR). There are three key UPR signal activators in mammals: IRE1α (inositol-requiring enzyme one alpha), PERK (protein kinase R-like ER kinase), and ATF6 (activating transcription factor 6). Activation of the endonuclease activity of IRE1α catalyzes removal of an intron from *XBP1* mRNA, resulting in production of Xbp1s, a transcription factor that drives the expression of genes involved in restoring protein homeostasis, localization and lipid synthesis (Adams et al., 2019). IRE1α also triggers regulated IRE1-dependent decay (RIDD), leading to degradation of RNAs in a cell and stimulus specific manner to regulate functions including apoptosis, metabolism and inflammation (Abdullah & Ravanan, 2018). In the PERK pathway, PERK phosphorylates eIF2α to inhibit global translation and at the same time promote translation of the ATF4 transcription factor. In the ATF6 pathway, ATF6 translocates to the Golgi network where is it cleaved by site 1 protease (S1P) to release the ATF6p50 transcription factor. ATF4 and ATF6p50 control the expression of an overlapping set of genes, including ones encoding proteins that contribute to protein folding in the ER. Activation of these pathways was initially ascribed to dissociation of BiP/GRP78 from each sensor, however it is now apparent that other ER chaperones are involved in a cell- and tissue-specific manner (Kettel & Karagöz, 2024).

UPR signaling events, and in particular the Ire1α pathway, are involved in the differentiation of many cell types, including myoblasts (He et al., 2021), fibroblasts (Baek et al., 2012), osteoblasts (Saito et al., 2011), neutrophils (Tanimura et al., 2018), plasma cells (Reimold et al., 2001), mammary epithelial cells (Tsuchiya et al., 2017), eosinophils (Bettigole et al., 2015), dendritic cells and macrophages (Bettigole & Glimcher, 2016). The UPR also has a vital role in sustaining the activity of specialized secretory cells such as plasma cells and pancreatic β cells (Hetz & Papa, 2018).

The role of UPR signaling in megakaryopoiesis and thrombopoiesis is unclear. It has been proposed that ER stress activation triggers apoptotic-like caspase activation during megakaryocytic differentiation of human megakaryoblastic cell lines, and may drive proplatelet formation (Kovuru et al., 2020; Lopez et al., 2013; Morishima & Nakanishi, 2016). However, while early reports suggested that localized caspase activation is required for megakaryopoiesis (Clarke et al., 2003; De Botton et al., 2002), the role of caspase activation in megakaryopoiesis is uncertain. Indeed genetic studies in mice convincingly showing that intrinsic apoptosis must be suppressed for successful differentiation (Josefsson et al., 2014; Josefsson et al., 2020) (McArthur et al., 2018). The studies suggesting a role for ER stress-induced apoptosis in megakaryopoiesis have mostly used PMA-induced maturation of various cell lines. However, the PMA concentrations used to induce maturation and draw conclusions about the role of UPR signaling also resulted in extensive cell death (Kovuru et al., 2020; Lopez et al., 2013; Morishima & Nakanishi, 2016). Therefore, it is unclear whether UPR activation in these studies was a specific feature of megakaryocytic maturation or a generalized response to cell stress. Here we investigate the involvement of UPR signaling pathways during in vitro maturation of megakaryocytic cell lines and human peripheral blood (PB) derived progenitors under conditions that induce maturation in the absence of cell death.

## Results

### The unfolded protein response is activated during megakaryocytic maturation of K-562 and MEG-01 cells

The K-562 and MEG-01 cell lines undergo megakaryocytic maturation upon PMA treatment (Herrera et al., 1998; Kamal et al., 2018). Previous studies of the ER stress response in various megakaryocytic cell lines have used concentrations of PMA (10-40 nM) that cause high levels (>30%) of cell death (Kovuru et al., 2020; Lopez et al., 2013). We confirmed that treatment with 0.5 nM PMA stimulated maturation as assessed by an increase in the expression of the transcription factor *GATA1* (Figure S1A&B) and the cell surface protein CD61 (Figure S1C). Importantly there was no change in cell viability (Figure S1D) or caspase activation (Table S1) during maturation, providing an experimental system in which we could study the role of the UPR in megakaryocytic maturation in the absence of confounding cell death/apoptosis.

To determine whether the UPR is activated during PMA-induced maturation of K-562 and MEG-01 cells we examined the expression of UPR target genes (Figure 1A and B). Expression of the IRE1α target *ERdj4* (*DNAJB9*) and the ATF6 targets *HERPUD1* and *XBP1* increased in response to PMA. The expression of *CHOP* (*DDIT3*), which is induced in response to all three UPR pathways also increased, whereas the expression of the PERK targets *GADD34* and *ATF4* increased in K-562 but not Meg-01 cells. The protein expression of BiP/GRP78, a UPR sensor and a UPR target downstream of ATF6, was unchanged at day 3 (Figure 1C). These results suggest that the activation of UPR signaling pathways occurs during megakaryocytic maturation. Since these changes in expression occurred without a loss of cell viability, this most likely represents involvement of the adaptive rather than terminal UPR. To determine whether this result was a feature of PMA-induced maturation of the cell lines, or also occurred during TPO-induced megakaryocyte differentiation, we analyzed the expression of *ERdj4*, *HERPUD1* and *CHOP* at day 11 (sample 1) or day 15 (sample 2) following TPO treatment of CD45^+^ cells isolated from human peripheral blood (Figure 1D). All three genes showed increased expression compared to day 0, but insufficient samples were available to determine statistical significance.

**Figure 1.**
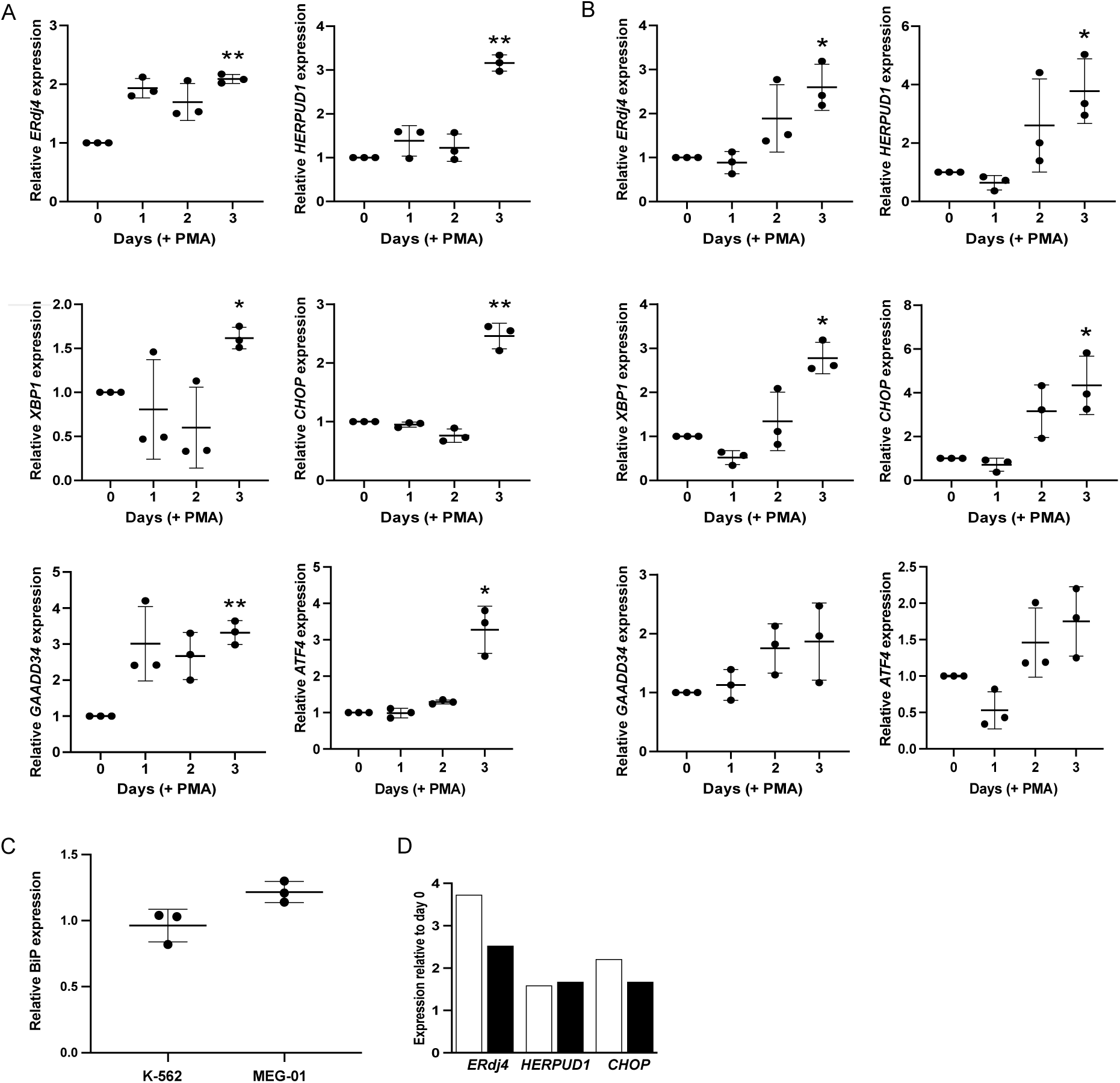
UPR target gene expression is upregulated during PMA-induced maturation of K-562 and MEG-01 cells into megakaryocytes. K-562 (A) or MEG-01 (B) cells were treated with 0.5 nM PMA and expression of *ERdj4, HERPUD1*, *XBP1, CHOP*, *GADD34* and ATF4 mRNA at days 0, 1, 2 and 3 was determined by RT-qPCR. Relative expression was calculated using the value at day 0 as 1. N=3 ± SD. \**P* < 0.05, \*\**P* < 0.01 compared to day 0 (one sample t-test). (C) K-562 or MEG-01 cells were treated with 0.5 nM PMA and BiP protein expression was determined by western blotting at day 3. Expression is relative to untreated cells. N=3 ± SD. (D) Expression (relative to day 0) of *ERdj4, HERPUD1* and *CHOP* following TPO-induced differentiation of CD45^+^ peripheral blood cells. White bars = sample 1 at day 11 of culture, black bars = sample 2 at day 15 of culture.

### Inhibition of IRE1α endonuclease activity, but not PERK or ATF6, inhibits megakaryocytic maturation

We next used inhibitors of the three UPR signaling pathways to determine whether activation of one or more contributes to megakaryocytic maturation. SN34221 inhibits the endonuclease activity of IRE1α (Patterson & Lonergan, 2008), AMG PERK 44 inhibits PERK (Smith et al., 2015), and PF-429242 inhibits the site-1 protease that cleaves ATF6 to release ATF6p50 (Hay et al., 2007). The inhibitors showed the expected specificity with SN34221 decreasing expression of *ERdj4*, AMG PERK 44 decreasing expression of *CHOP*, *HERPUD1* and possibly *GADD34*, and PF-429242 decreasing expression of *CHOP* and *HERPUD1* at day 3 of PMA-induced megakaryocytic maturation of K-562 cells (Figure S2). Inhibition of IRE1α had a negative impact on maturation, with 25% lower CD61 expression and 40% lower *GATA1* expression at day 3 at the highest concentration of the inhibitor (Figure 2A&D). Treatment with 20 and 50 µM SN34221 was associated with a small decrease in cell viability at day 3 (Figure S2D). In contrast when PERK was inhibited with AMG PERK 44, expression of CD61 increased, suggesting an enhancement of maturation, not *GATA1* expression was unchanged, suggesting no impact on maturation (Figure 2B&D). These conflicting results may have occurred because AMG PERK 44 also led to up to 50% loss in cell viability (Figure S2E), and while CD61 expression was assessed only in the live cells, *GATA1* expression was assessed in the total cell population. Thus the PERK pathway may be important in both preventing apoptosis of the maturing cells and in the maturation process. Finally, inhibition of the ATF6 pathway had no impact on maturation (Figure 2C&D) or cell viability (Figure S2F).

**Figure 2.**
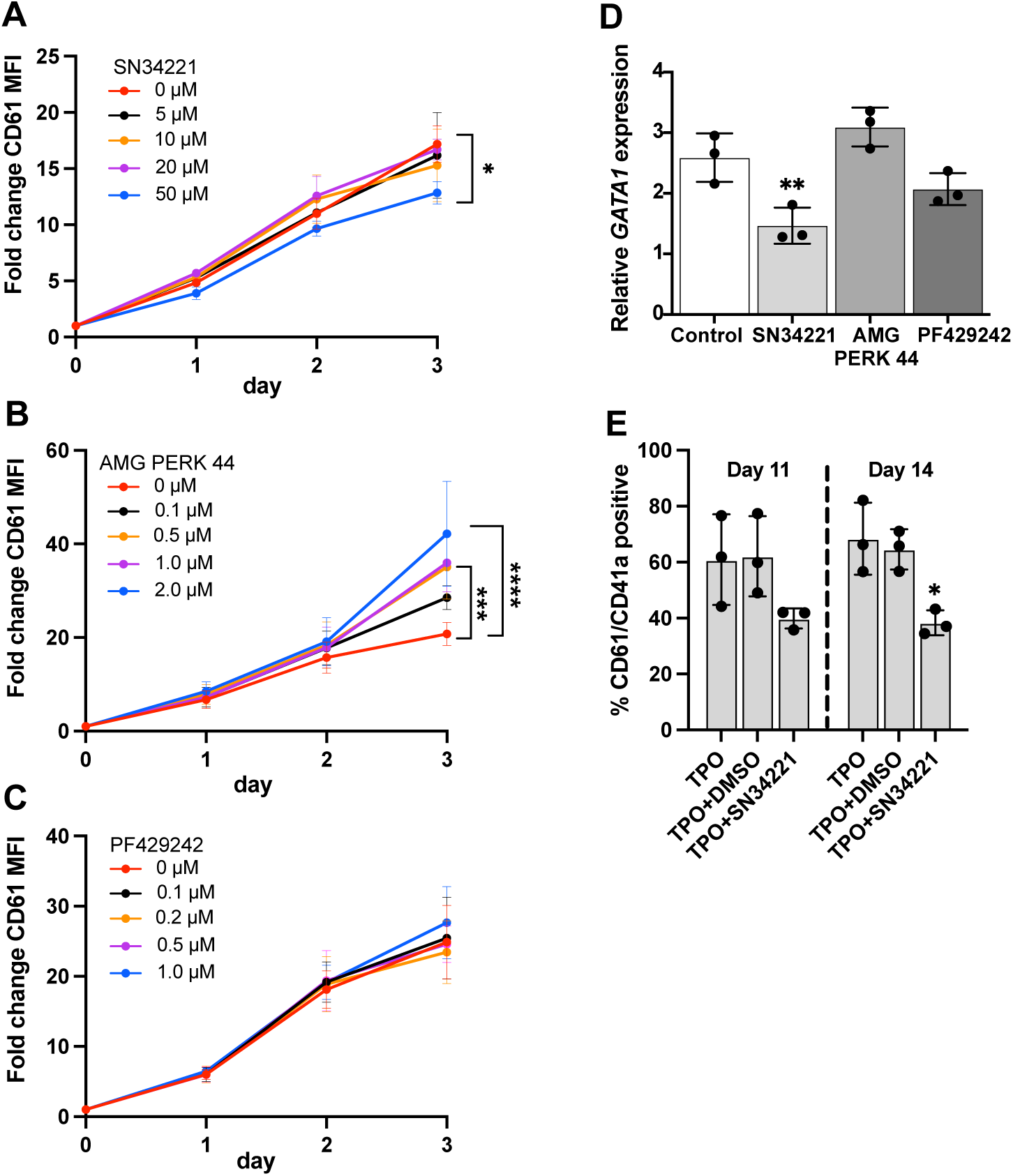
The effect of pharmacological inhibition of UPR on K-562 maturation and peripheral blood CD45^+^ cell differentiation. **A - C**. K-562 cells were treated with 0.5 nM PMA in the presence of SN34221 (A), AMG PERK 44 (B) of PF429242 (C) and expression of CD61determined (fold change MFI compared to day 0). N=3 ± SD \**P* < 0.05, \*\*\**P* < 0.001, \*\*\*\**P* < 0.0001 (two-way ANOVA). **D.** K-562 cells were treated with 0.5 nM PMA in the presence (white bars) or absence (grey bars) of 50 μM SN34221, 2 μM AMG PERK 44 or 1 μM PF429242 and relative *GATA1* expression measured at day 3. N=3 ± SD. \*\**P* < 0.01, (one-way ANOVA). **E.** % CD61^+^/CD41a^+^ cells at days 11 and 14 of TPO-induced megakaryocyte differentiation of CD45^+^ peripheral blood cells in the presence of DMSO or 50 μM SN34221. N=3 ± SD \**P* < 0.05 compared to TPO + DMSO (one-way ANOVA).

Having observed the potential involvement of the IRE1α UPR pathway in megakaryocytic maturation of K-562 cells, we next determined the impact of IRE1α inhibition on TPO-induced megakaryocyte differentiation of CD45^+^ cells isolated from human peripheral blood. The IRE1α endonuclease inhibitor SN34221 significantly decreased the percentage of CD61^+^/CD41a^+^ megakaryocytes in the live cell population at days 11 and 14 (Figure 2E). Consistent with this result, when cultures were examined by bright field microscopy the cells treated with SN34221 were generally smaller and there was no evidence of proplatelet formation or membrane blebbing (Figure S3C). Furthermore, we did not see an increase in the percentage of CD61^+^/CD41a^+^ megakaryocytes in the non-viable (Zombie stained) cells in the cultures. These results suggest that the IRE1α signaling branch of UPR may be involved in PMA-induced megakaryocytic maturation of K-562 cells and TPO-induced megakaryocyte differentiation of primary cells.

### Activation of UPR does not enhance PMA-induced maturation of K-562 cells

Our data suggests that activation of the UPR, in particular the IRE1α and PERK pathways, may be involved in megakaryocyte maturation. We next determined whether activation of UPR signaling by an exogenous ER stress enhanced megakaryocyte maturation. We confirmed that the ER stressors DTT, a small-molecule redox reagent for reducing protein disulfide bonds, and thapsigargin, a sarco/endoplasmic reticulum ATPase (SERCA) inhibitor, induced UPR as shown by the splicing of the IRE1α target *XBP1* (Figure S4). Treatment with DTT had no impact on megakaryocytic maturation (Figure 3A) or viability (Figure 3B) of K-562 cells. In contrast, treatment with thapsigargin caused a dose-dependent reduction in PMA-induced maturation of K-562 cells (Figure 3C) with no change in cell viability (Figure 3D). Thapsigargin also inhibited TPO-induced differentiation of CD45^+^ peripheral blood cells (Figure 3E) and subsequent budding and proplatelet production (Figure S3D). The difference in effect of the two ER stressors suggests that the exogenous UPR activation does not enhance megakaryocyte maturation, whereas alteration of ER Ca^2+^ homeostasis inhibits megakaryocyte differentiation.

**Figure 3.**
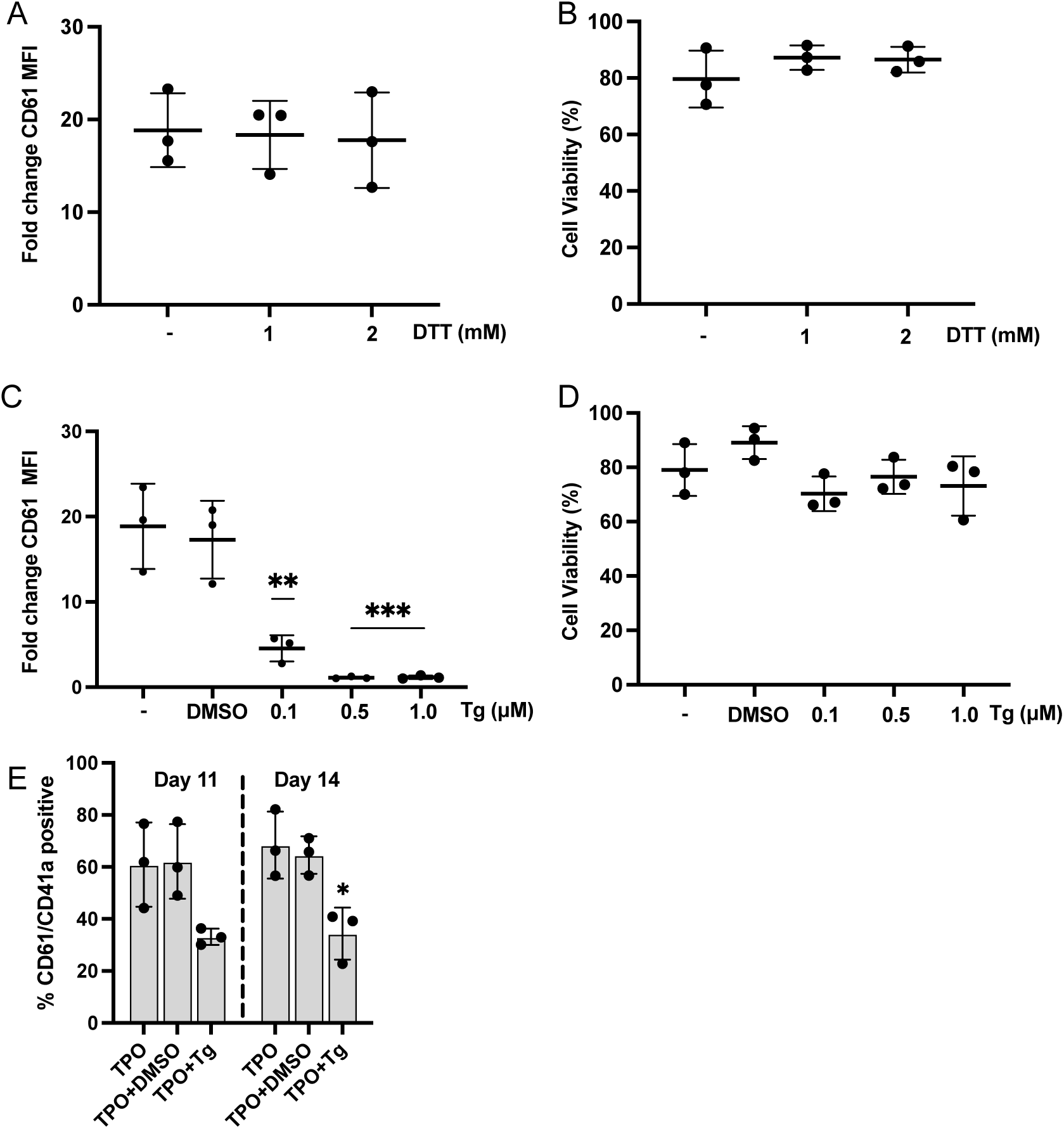
The effect of UPR activation on K-562 maturation and peripheral blood CD45^+^ cell. **A - D** K-562 cells were treated with 0.5 nM PMA for 2 days in the presence of DTT (A and B) or Tg (C and D) and analyzed for expression of CD61 (fold change MFI compared to day 0) (A and C) or cell viability (B and D). N=3 ± SD. \**P* < 0.05, \*\**P* < 0.01 (one way ANOVA). **E.** % CD61^+^/CD41a^+^ cells at days 11 and 14 of TPO-induced megakaryocyte differentiation of CD45^+^ peripheral blood cells in the presence of DMSO or 1 μM Tg. N=3 ± SD \**P* < 0.05 compared to TPO + DMSO (one-way ANOVA).

### Megakaryocytic maturation of K-562 or MEG-01 cells does not alter the response to exogenous ER stressors

Finally, we examined whether maturation alters the ability of the K-562 or MEG-01 cells to respond to exogenous ER stressors. K-562 cells or MEG-01 cells were treated with DTT or thapsigargin before or 3 days after PMA treatment, and the expression of UPR target genes was quantified by RT-qPCR. DTT treatment activated the UPR as indicated by a strong induction of *XBP1* splicing, and *CHOP* and *BIP* expression, in immature (untreated) and mature (PMA treated) K-562 cells (Figure 4A). Similarly, thapsigargin treatment induced *XBP1* splicing and increased *CHOP, BIP* and *GADD34* expression in immature and mature K-562 cells (Figure 4B). Similar results were obtained with MEG-01 cells where both ER stressors activated UPR signaling in immature and mature cells (Figure S5). These results indicate that even though UPR pathways are activated during PMA-induced maturation, this does not alter the ability of the cells to further activate UPR signaling in response to exogenous ER stressors.

**Figure 4.**
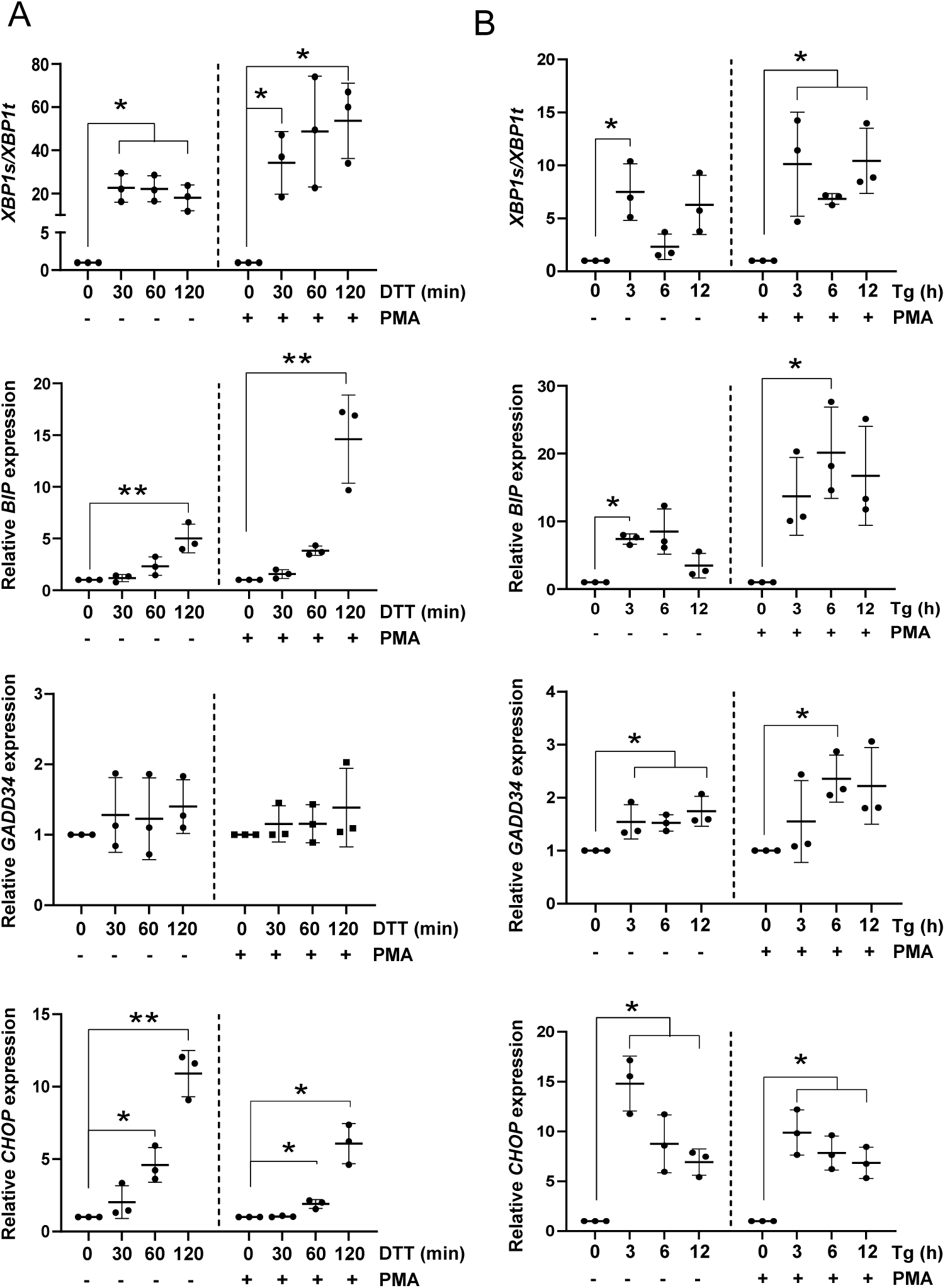
PMA treated and untreated K-562 cells are similarly responsive to ER stress induced by DTT or thapsigargin (Tg) Immature (no PMA) or mature (3 days after PMA treatment) K-562 cells were treated with 10 mM DTT for 0-2 h (A) or 0.1 µM Tg for 0-12 h (B). Expression of *XBP1*s and *XBP1*t, *BIP, CHOP* and *GADD34* mRNA was quantified by RT-qPCR. Expression is relative to 0 h. N=3 ± SD. \**P* < 0.05, \*\**P* < 0.01 (one sample t-test).

## Discussion

In the current study we used a low concentration of PMA that induced megakaryocytic maturation of K-562 and MEG-01 cell lines in the absence of cell death. Under these conditions maturation was accompanied by increased transcription of UPR target genes *ERDj4, HERPUD1*, *XBP1*, *CHOP*, and *GADD34,* indicating activation of all three UPR signaling pathways. These results are consistent with previous studies reporting an increase in expression of UPR target genes in various models of megakaryocytic maturation (Kovuru et al., 2020; Lopez et al., 2013; Morishima & Nakanishi, 2016). However, in contrast to these earlier studies, increased expression of UPR target genes occurred in the absence of cell death, demonstrating that UPR-induced apoptosis is not required for megakaryocytic maturation. Increased expression of *ERdj4*, *HERPUD* and *CHOP* also occurred during TPO-induced differentiation of CD45^+^ cells, and an increase in BiP/GRP78 in differentiating CD34^+^ cells has been reported (Lopez et al., 2013). The terminal stages of megakaryopoiesis involve endomitosis and polyploidization with an accompanying high demand for synthesis of proteins and membrane expansion and remodeling prior to proplatelet formation and platelet release (de Jonckheere et al., 2023; Mazzi et al., 2018). Therefore, it is unsurprising that the adaptive UPR is activated in maturing megakaryocytes to assist terminal megakaryopoiesis by increasing the biosynthesis of chaperone proteins, increasing membrane synthesis, and boosting the protein folding capacity of the ER.

To determine whether activation of a specific arm of the adaptive UPR was particularly important for megakaryopoiesis, we inhibited IRE1α endonuclease activity (SN34221), PERK autophosphorylation (AMG PERK 44), and ATF6 cleavage (PF-429242). The IRE1α inhibitor SN34221 partially inhibited PMA-induced maturation of K-562 cells and TPO-induced differentiation of peripheral blood CD45^+^ cells, implicating the IRE1α UPR pathway in supporting megakaryocytic maturation. Whether *XBP1* splicing and/or RIDD activation is relevant for human megakaryopoiesis remains an open question and requires further investigation. XBP1s is involved in maturation of other hematopoietic cell types (Bettigole & Glimcher, 2016; Reimold et al., 2001), and is an important regulator of specialist secretory cells which require a high level of protein synthesis (Lee et al., 2005; Reimold et al., 2001). XBP1s regulates gene expression by forming heterodimers with other transcription factors including MIST1, an important regulator of differentiation (Acosta-Alvear et al., 2007; Hetz & Papa, 2018). A recent characterization of mice with megakaryocyte/platelet-specific deletion of *XBP1* found normal platelet counts but a higher percentage of circulating reticulated (newly released) platelets (Jain et al., 2022) suggesting possible alteration in megakaryopoiesis and/or thrombopoiesis. RIDD has also been implicated in controlling differentiation-associated processes (He et al., 2021; Levi-Ferber et al., 2021) but has not been directly studied in the context of megakaryocyte biology. In contrast, inhibition of the PERK pathway enhanced maturation but also increased cell death, suggesting that this pathway is critical for preventing megakaryocyte apoptosis, but may act to restrain megakaryocyte maturation. This result is consistent with inhibition of PMA-induced megakaryocytic maturation by salubrinal, a drug which potentiates eIF2α phosphorylation (Boyce et al., 2005; Lopez et al., 2013), but in contrast to the decreased platelet count seen in mice with megakaryocyte/platelet-specific deletion of *PERK* (Jain et al., 2022). However, the high level of cell death we saw with PERK inhibition may explain the decreased platelet count observed by Jain et al (2022).

Our observations that both immature and mature K-562 and MEG-01cells show similar responses to the exogenous ER stressors DTT and thapsigargin, and that DTT has no impact on megakaryocytic maturation, suggest that the mechanisms of action of the IRE1α and PERK pathways are distinct from a canonical ER stress response. Further studies, including with additional inhibitors, are needed to tease out which downstream targets of XBP1s/RIDD/ATF4 are involved in human megakaryopoiesis and thrombopoiesis, and to delineate the role of the UPR pathways in preventing cell death versus facilitating differentiation. It would also be interesting to determine the impact of tunicamycin, which in platelets specifically activates the IRE1α pathway (Jain et al., 2022), on TPO-induced CD45^+^ or CD34^+^ differentiation.

Treatment with thapsigargin inhibited both PMA-induced maturation of K-562 cells and TPO-induced differentiation of peripheral blood-derived CD45^+^ cells. Thapsigargin blocks SERCA, inducing protein misfolding and the UPR, and altering Ca^2+^ homeostasis (Carreras-Sureda et al., 2018; Jaskulska et al., 2020; Sehgal et al., 2017). Since DTT had no impact on megakaryocytic maturation, it is likely that dysregulation of Ca^2+^ homeostasis underlies inhibition of maturation by thapsigargin. This is consistent with the known roles of intracellular Ca^2+^ in megakaryopoiesis. Activation of phospholipase C generates inositol trisphosphate (IP3) triggering release of Ca^2+^ from the ER via the IP3 receptor. Subsequent store-operated calcium entry (SOCE) induces expression of nuclear factor of activated T cells (NFAT), a transcriptional regulator implicated in megakaryopoiesis (Hogan et al., 2003; Zaslavsky et al., 2013; Zhu et al., 2018). Additionally, Ca^2+^entry through a membranous Ca^2+^ ion channel N-methyl-D-aspartate receptor supports megakaryocyte differentiation (Hearn et al., 2020; Hearn et al., 2022). Ca^2+^-dependent activation of Ca^2+^/calmodulin-dependent protein kinase (CaMK) induces transcription of CREB (c-AMP response element-binding protein) to activate the genes with CRE elements, including megakaryocyte-lineage specific genes such as *FLI1*, *HOXC6*, *MXD3*, *PRDM16*, *FOXB1*, *TBX6*, *HDAC11* and *NPAS1* (Hogan et al., 2003; Zhang et al., 2005; Zhu et al., 2018). Finally, changes in intracellular Ca^2+^ homeostasis occur in myeloproliferative neoplasms and may contribute to disease pathogenesis through ER stress (Di Buduo et al., 2020; Immanuel et al., 2022; Pietra et al., 2016; Seetharam et al., 2023). Thus, our data provide further evidence for the importance of calcium signaling in megakaryopoiesis.

Megakaryopoiesis and thrombopoiesis are complex process that require the activation of many synergistic pathways. Overall, our findings indicate that the adaptive UPR, rather than terminal UPR and associated apoptosis, is a feature of megakaryocyte differentiation. Our data also provide preliminary evidence that activation of the IRE1α pathway enhances maturation, whereas the PERK pathway restrains maturation. We speculate that the two pathways may be active at different times to support different phases of differentiation. To understand the interplay between activation of these two pathways, future work should focus on determining the nature of the initiating events (e.g. is BiP and/or another chaperone important), and the timing of IRE1α phosphorylation and PERK phosphorylation, during differentiation.

### Experimental Procedures

#### Cell culture and differentiation

Human myeloid leukemic cells, MEG-01 (ATCC CRL-2021) (Ogura et al., 1985) and K-562 (ATCC CRL-243) (Tabilio et al., 1983), were maintained in RPMI-1640 (Gibco; 3180002) medium supplemented with 10% fetal bovine serum (FBS), sodium bicarbonate (2 g/L), penicillin (100 U/mL), and streptomycin (100 µg/mL) at 37°C in a 5% CO_2_ humidified atmosphere. The identity of the cell lines was confirmed by STR analysis (DNA Diagnostics, NZ) and the cells were confirmed to be mycoplasma free. In vitro megakaryocytic maturation was induced by treating K-562 or MEG-01 cells (2.5 x 10^5^ cells/mL) in a 6-well plate with 0.5 nM phorbol-12-myristate-13-acetate (PMA) (Sigma-Aldrich).

CD45^+^ cells were isolated and purified from peripheral blood mononuclear cells from individual donors as previously described (Balduini et al., 2009; Ong et al., 2017). To induce progenitor differentiation into megakaryocytes, isolated CD45^+^ cells (1 x 10^5^ cells/ml) were cultured in StemSpan^TM^ medium (StemCell Technologies, Vancouver, Canada) supplemented with 10 ng/ml thrombopoietin (TPO), interleukin (IL) 6 and IL 11 (StemCell Technologies). Cultures were maintained at 37°C in a 5% CO_2_ humidified atmosphere for 14 days with media changes on days 5, 8, and 11.

#### ER stress modulators

The conventional ER stressors thapsigargin (Tg) (T9033; Sigma-Aldrich) or dithiothreitol (DTT) (D0632; Sigma-Aldrich) were used at the indicated concentrations and time points. The following compounds were used as UPR inhibitors: PF429242-dihydrochloride (SML0667; Sigma-Aldrich), an ATF6 inhibitor (Hay et al., 2007), AMG-PERK-44 (HY-12661A, MedChemExpress), a PERK inhibitor (Smith et al., 2015), and SN34221 (Supplementary data), an IRE-1α endonuclease inhibitor (Patterson & Lonergan, 2008). The chemicals were dissolved in DMSO or distilled water, and DMSO used as a control when appropriate.

#### Analysis of megakaryocytic maturation by flow cytometry

Cells were harvested, washed with Dulbecco’s PBS, resuspended in 100 µL of Dulbecco’s PBS, and stained for 30 min in the dark at room temperature. Antibodies used were anti-CD61 phycoerythrin (PE) (1:20; 1075384; BD Biosciences), mouse IgG1 PE (isotype control; 7223589; BD Biosciences), anti-CD41a V450 (1:20; 1011309; BD Biosciences), V450 IgG1 (isotype control; 7128864; BD Biosciences). For K-562 and MEG-01, live cells were determined with Zombie NIR (1:100; B312060; BioLegend) or Zombie Green (1:100; B334686; BioLegend). After incubation, the cells were washed with Dulbecco’s PBS or wash buffer (1X Dulbecco’s PBS, 1 mM EDTA, 3% FBS), resuspended in FACS buffer (0.01% sodium azide, 0.1% BSA in Dulbecco’s PBS) analyzed using the Guava® easyCyte 5 HPL benchtop flow cytometer (MerckMillipore) or BD LSRFortessa^TM^ cell analyzer (BD Biosciences). Data analysis was performed using FlowJo version 10 (FlowJo, Ashland, USA). Data reported are geometric mean fluorescence intensity (MFI) of the Zombie negative population.

#### Caspase assay

K-562 cells were seeded into 96-well plates at 2.5 x 10^5^ cells/mL in 6-well plate and treated with 0.5 nM PMA or 10 mM DTT as a positive control. At 1, 2 or 3 days (PMA) or 22 h (DTT) cells were washed with PBS, harvested and resuspended in100 μL caspase assay buffer (50 mM Hepes pH 7.25, 10% sucrose, 5 mM DTT, 0.1 % CHAPS). Following the addition of 50 μM Ac-DEVD-AMC the production of AMC (λ_EM_ = 390 nm, λ_EX_ = 460 nm) was assayed over 60 min at 37 °C in a CLARIOstar_™_ microplate reader (BMG LABTECH). The results are expressed as fold increase in activity compared to untreated.

#### RNA isolation and RT-qPCR

Total RNA was isolated from K-562, MEG-01 and primary cells using TRIzol (Invitrogen). The concentration of the RNA was measured using Qubit^TM^ RNA HS Assay Kit (ThermoFisher Scientific). First-strand cDNA synthesis was performed with 1 µg of total RNA in 20 µL reaction volume using SuperScript^TM^ IV VILO^TM^ (SSIV VILO) master mix. Real-time quantitative RT-PCR reactions were run on LightCycler^TM^ 480 Instrument II System (Roche Life Science) using PowerUp^TM^ SYBER^TM^ Green Master Mix (Thermo Fisher Scientific). All the procedures were followed as recommended by the manufacturer. The qPCR primer sequences are shown in Table S2. RT-qPCR was performed in triplicate, and relative expression was quantified using the Pfaffl method (Pfaffl, 2001). Data were normalized to *HPRT* and *GAPDH* (K-562 and MEG-01 cells) or *RPL27* (primary cells) using basket normalization (Vandesompele et al., 2002).

#### Semi-quantitative PCR analysis for XBP1 splicing detection

To detect *XBP1* splicing, *XBP1* mRNA was amplified (forward primer: 5′ CCTTGTAGTTGAGAACCAGG 3′; reverse primer 5′ AGGGGCTTGGTATATATGTGG 3′) (Fribley et al., 2011). Briefly, first-strand cDNA was synthesized with 2 µg of total RNA in 20 µL reaction volume. To amplify *XBP1* 5 µL of undiluted cDNA was used as a template in a 50 µL reaction mixture with an initial denaturation step for 3 min at 94 °C, followed by 30 cycles of denaturation for 30 s at 94 °C, annealing for 30 s at 54 °C, and extension for 60 s at 72 °C, and a final extension step for 10 min at 72 °C. PCR products were resolved on a 2% (w/v) agarose gel, stained with 0.5 µg/mL ethidium bromide, and visualized using the Gel Doc^TM^ XR+ Imaging System (Bio-Rad).

#### Protein extraction and western blotting

Cells were lysed in RIPA (radio-immunoprecipitation assay) buffer (50 mM Tris-HCl pH 7.4, 150 mM NaCl, 1 mM EDTA, 1% NP-40, 0.25% sodium deoxycholate) containing protease and phosphatase inhibitor cocktail (Sigma-Aldrich; MSSAFE). Total protein (20 µg) was resolved by SDS PAGE under reducing conditions and transferred to nitrocellulose membrane. Membranes were stained with REVERT^TM^ 700 total protein stain (LI-COR Biosciences), washed and imaged on the Odyssey Fc Imaging System (LI-COR Biosciences). The membrane was then blocked with Intercept® Blocking Buffer (LI-COR Biosciences) for 1 h at room temperature and incubated overnight with 1:500 anti-GRp78 (BiP) antibody (sc-1050; Santa Cruz) at 4 °C. IRDye® 680-LT donkey anti-goat (C60127-06, LI-COR Biosciences) was used as a secondary antibody, and the blots were visualized in the Odyssey Fc Imaging System. Band intensities were quantified with Image Studio Lite version 5.2 (LI-COR Biosciences). Briefly, signals from the total protein stain and target protein were measured, the lane normalization factor (LNF) was calculated for total protein in each sample, and the target protein band intensities were normalized using the LNF.

#### Statistical analysis

Statistical analyses were performed using GraphPad Prism software version 8.0 (GraphPad Software, San Diego, CA, USA). The comparison of means between different groups was performed using one sample t-test, one-way or two-way ANOVA with Dunnett’s multiple comparisons test, or Mann-Whitney. *P* < 0.05 was considered significant. For ANOVA, normality of residuals was confirmed using the D’Agostino, Anderson-Darling, Shapiro-Wilk and Kolmogorov-Smirnov tests. For all experiments the number of biological replicates is indicated.

## Supplementary Materials

The supplementary data file contains the following:

Synthesis of SN34221

Figure S1. PMA induces megakaryocyte maturation of K-562 and MEG-01 cells

Figure S2. Impact of UPR inhibitors on cell viability and maturation of K-562 cells

Figure S3. Impact of SN34221 or thapsigargin on proplatelet production

Figure S4. Thapsigargin and DTT induce *XBP1* splicing in K-562 cells

Figure S5. UPR activation in immature and mature MEG-01 cells in response to DTT or thapsigargin

Figure S6. SN34221 inhibits *XBP1* splicing.

Table S1. Caspase activation during PMA-induced maturation of K-562 cells

Table S2. RT-qPCR primers used in the study

## Statements and Declarations

### Author contributions

ECL and MK-Z conceived the study. ECL, MK-Z and MF designed the study. Experimental work was performed by MF and CD-H. All authors contributed to data analysis. The first draft of the manuscript was written by MF and all authors commented on previous versions of the manuscript. All authors read and approved the final manuscript.

### Data availability

Data and materials included within the article or Supplementary Materials are available from the authors on request.

### Preprint

This article has been deposited on BioRxiv: https://www.biorxiv.org/content/10.1101/2023.07.26.550730v1.full

### Ethics approval

The study was approved by the University of Otago Human Ethics Committee (NZ) (H20/132).

### Consent to participate

Written informed consent was obtained from all individual participants included in the study.

