## Supplementary Data for "Megakaryocyte maturation involves activation of the adaptive unfolded protein response"

Synthesis of SN34221	page 2

Figure S1. PMA induces megakaryocyte maturation of K-562 and MEG-01 cells.	page 4

Figure S2. Impact of UPR inhibitors on cell viability and maturation of K-562 cells.	page 5

Figure S3. Impact of SN34221 or thapsigargin on proplatelet production.	page 6

Figure S4. Thapsigargin and DTT induce *XBP1* splicing in K-562 cells.	page 7

Figure S5. UPR activation in immature and mature MEG-01 cells in response to DTT or thapsigargin.	page 8

Figure S6. SN34221 inhibits *XBP1* splicing.	page 9

Table S1. Caspase activation during PMA-induced maturation of K-562 cells	page 10

Table S2. RT-qPCR primers used in the study.	page 11

**Synthesis of SN34221 (4).** The synthesis of SN34221 was carried out using methodology previously described^1^. SN34221 was described as having an IC_50_ for inhibition of cleavage of an IRE-1α mini human XBP1 mRNA stem loop substrate of 3 nM, and a cellular EC_50_ for splicing of *XBP1* of 5 µM^1^. In K-562 cells the cellular EC50 for DTT-induced splicing of *XBP1* was ~17 μM (Figure S6).

**General procedures.** Final product purity was analyzed by reverse-phase HPLC, (Altima C18 5μm column, 3.2 × 150 mm) using an Agilent Technologies 1260 Infinity equipped with a diode-array detector. Mobile phases were gradients of 80% acetonitrile/20% H2O (v/v) in 45 mM ammonium formate at pH 3.5 and 0.7-1.2 mL/min. Final compound purity was determined by monitoring at 330 ± 50 nM. Melting points were determined on an Electrothermal 2300 Melting Point Apparatus. NMR spectra were obtained on a Bruker Avance 400 spectrometer at 400 MHz for 1H spectra. Chemical shifts and coupling constants were recorded in units of ppm and Hz, respectively. Low resolution mass spectra were gathered by direct injection of methanolic solutions into an Agilent 6120 mass spectrometer using an atmospheric pressure chemical ionization (APCI) mode with a fragmentor voltage of 50 V and a drying gas temperature of 250 °C. Organic solutions were dried over Na_2_SO_4_ and solvents were evaporated under reduced pressure on a rotary evaporator. Thin-layer chromatography was carried out on aluminium-backed silica gel plates (Merck 60 F254) with visualization of components by UV light (254 nm) or exposure to I_2_. Column chromatography was carried out on silica gel (Merck 230–400 mesh). DCM refers to dichloromethane, DMF refers to dimethylformamide, DMSO dimethyl sulfoxide, EtOAc refers to ethyl acetate, MeOH refers to methanol, pet. ether refers to petroleum ether boiling fraction 40–60 °C.

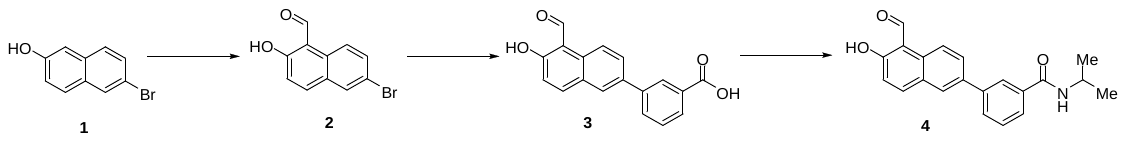

**6-Bromo-2-hydroxy-1-naphthalenecarbaldehyde (2).** A solution of TiCl_4_ (5.2 mL, 47.0 mmol) and dichloromethyl methyl ether (2.23 mL, 24.6 mmol) in dry DCM (10 mL) was stirred at 0 °C for 15 min. A suspension of 6-bromo-2-napthol (**1**) (5.0 g, 22.4 mmol) in dry DCM (50 mL) was added dropwise and the solution allowed to warm to 20 °C and stirred for 12 h. 1 M HCl (100 mL) was added and the mixture extracted with DCM (3 × 50 mL). The organic fraction was washed with water (50 mL) and dried and the solvent evaporated to give aldehyde **2** (3.39 g, 60%) as a white powder: mp (DCM) 139–141 °C; ^1^H NMR (CDCl_3_) δ 13.11 (s, 1 H, OH), 10.78 (s, 1 H, CHO), 8.22 (d, *J* = 9.0 Hz, 1 H, H-8), 7.96 (d, *J* = 2.1 Hz, 1 H, H-5), 7.89 (d, *J* = 9.1 Hz, 1 H, H-4), 7.69 (dd, *J* = 9.1, 2.1 Hz, 1 H, H-7), 7.18 (d, *J* = 9.1 Hz, 1 H, H-3); MS *m/z* 265.5, 267.5 (MH^+^, 100%).

**3-(5-Formyl-6-hydroxy-2-naphthalenyl)benzoic Acid (3).** Pd(PPh_3_)_4_ (272 mg, 0.24 mmol) was added to a stirred, N_2_-purged mixture of bromide **2** (1.18 g, 4.7 mmol), 3-carboxyphenylboronic acid (0.86 g, 5.17 mmol) and Na_2_CO_3_ (3.0 g, 28.2 mmol) in DMF/water (1:1, 50 mL) and the mixture was stirred at 105 °C for 5 h. The mixture was cooled to 20 °C and 1 M NaOH (30 mL) solution was added. The mixture was extracted with DCM (3 × 20 mL). The aqueous fraction was acidified with 6 M HCl solution, and stirred at 0 °C for 30 min. The precipitate was filtered, washed with water (10 mL), washed with Et_2_O (5 mL) and dried. The solid was suspended in hot MeOH/EtOAc (1:1, 50 mL) and filtered to give the acid **3** (0.96 g, 70%) as a white solid: ^1^H NMR [(CD_3_)_2_SO] δ 13.13 (s, 1 H, OH), 11.99 (br s, 1 H, CO_2_H), 10.83 (s, 1 H, CHO), 9.06 (d, *J* = 8.9 Hz, 1 H, H-4′), 8.83 (t, *J* = 1.6 Hz, 1 H, H-2), 8.25–8.29 (m, 2 H, H-1′, H-8′), 8.06 (ddd, *J* = 7.7, 1.8, 1.2 Hz, 1 H, H-6), 8.00 (dd, *J* = 8.9, 2.1 Hz, 1 H, H-3′), 7.96 (dt, *J* = 7.8, 1.2 Hz, 1 H, H-4), 7.64 (t, *J* = 7.8 Hz, 1 H, H-5), 7.30 (d, *J* = 9.0 Hz, 1 H, H-7′); MS *m/z* 291.3 (M-H, 100%).

**3-(5-Formyl-6-hydroxy-2-naphthalenyl)-*N*-(1-isopropyl)benzamide (4).** HBTU (169 mg, 0.45 mmol) was added to a stirred solution of acid **3** (100 mg, 0.34 mmol) and iPrNEt (118 μL, 0.68 mmol) in dry DMF (5 mL) and the mixture was stirred at 20 °C for 10 min. iPrNH_2_ (116 μL, 1.36 mmol) was added and the mixture was stirred at 20 °C for 2 h. The mixture was diluted with EtOAc (50 mL) and washed with water (3 × 10 mL). The solvent was evaporated and the residue was dissolved in 2 M HCl/MeOH (20 mL) and stirred for 16 h. The solvent was evaporated and the crude solid was purified by column chromatography, eluting with a gradient (50–100%) of EtOAc/pet. ether, to give benzamide **4** (89 mg, 70%) as a white powder: mp (EtOAc) 250–253 °C; ^1^H NMR [(CD_3_)_2_SO] δ 11.98 (br s, 1 H, OH), 10.84 (s, 1 H, CHO), 9.06 (d, *J* = 8.9 Hz, 1 H, H-4′), 8.36 (br d, *J* = 7.7 Hz, 1 H, CONH), 8.20–8.29 (m, 3 H, H-2, H-1′, H-8′), 8.03 (dd, *J* = 8.9, 2.0 Hz, 1 H, H-3′), 7.94 (dd, *J* = 7.8, 1.6 Hz, 1 H, H-4), 7.94 (dd, *J* = 7.8, 1.6 Hz, 1 H, H-4), 7.85 (dd, *J* = 7.8, 1.6 Hz, 1 H, H-6), 7.58 (t, *J* = 7.8 Hz, 1 H, H-5), 7.30 (d, *J* = 9.0 Hz, 1 H, H-7′), 4.10–4.18 (m, 1 H, CHN), 1.20 (d, *J* = 6.6 Hz, 6 H, 2 × CH_3_); MS *m/z* 334.6 (MH^+^, 100%). Anal. calcd for C_21_H_19_NO_3_·¾H_2_O: C, 72.71; H, 5.96; N, 4.04. Found: C, 73.14; H, 6.07; N, 3.73%. HPLC purity 98.0%.

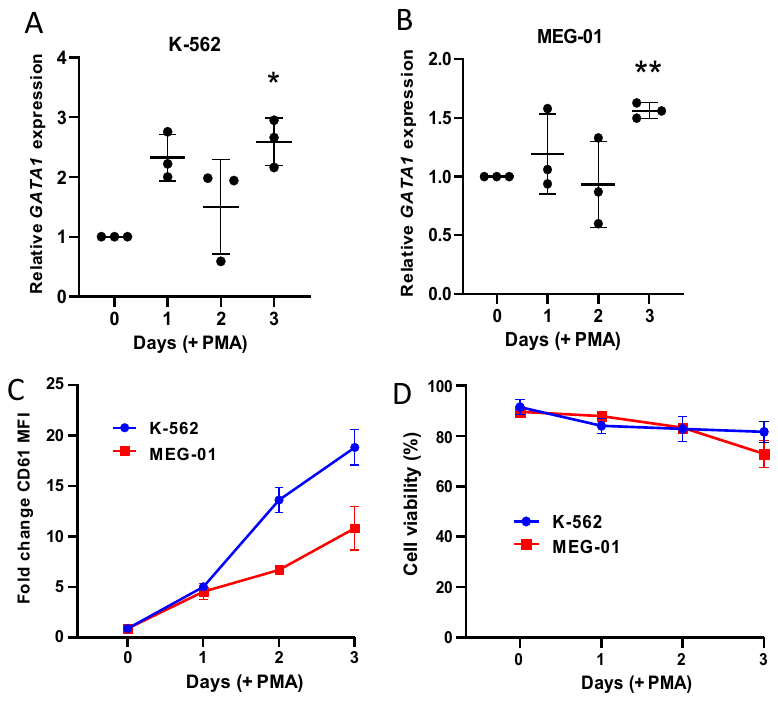

**Figure S1.** **PMA induces megakaryocyte maturation of K-562 and MEG-01 cells.** Cells were treated with 0.5 nM PMA and analyzed at days 0, 1, 2 and 3. **A** and **B.** RT-qPCR analysis of *GATA1* mRNA expression for K-562 (A) and MEG-01 (B) cells. Relative expression is calculated using the value at day 0 as 1. N = 3 ± SD. **P* < 0.05, ***P* < 0.01 (one sample t-test). **C.** Quantification of PE-CD61 (fold change in MFI compared to day 0). **D.** Cell viability (% Zombie negative).

**Figure S2.** **Impact of UPR inhibitors on UPR gene expression and cell viability. A-C.** Expression of indicated genes at day 3 of PMA-induced maturation in the absence (white bars) or presence (grey bars) of 50 μM SN4221 (A), 2 μM AMG PERK 44 (B) or 1 μM PF429242 (C). N = 3 ± SD. **P* < 0.05 compared to DMSO control (Mann-Whitney). **D-F**. Flow cytometry analysis of cell viability using Zombie Green at days 0-3 in K-562 cells treated with SN4221 (D), AMG PERK 44 (E) or PF429242 (F). N = 3 ± SD. There was a significant decrease in cell viability compared to 0 h for 20 µM and 50 µM SN4221 (day 3), and all concentrations of AMG PERK 44 (days 2 and 3) (two-way ANOVA).

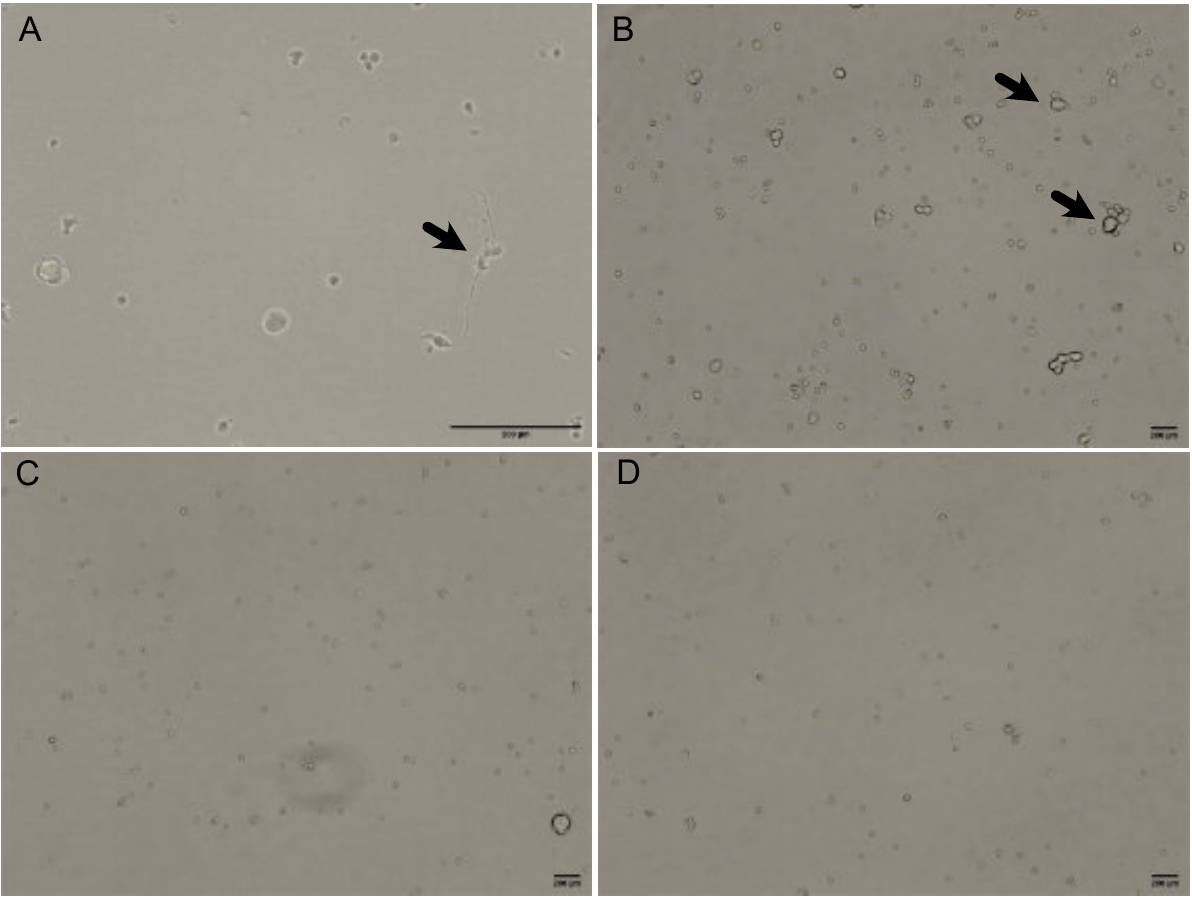

**Figure S3. Impact of SN34221 or thapsigargin on proplatelet production.** Bright field images of cellular morphology during TPO-induced differentiation of human peripheral blood CD45+ cells at day 11 in either untreated cells (A and B) or cells treated with 50 μM SN34221 (C) 2 μM Tg (D). Scale bars are 200 μm. Arrows indicate proplatelet forming (A) and budding (B) cells. Images were taken using the EVOS XL Core imaging system (ThermoFisher Scientific).

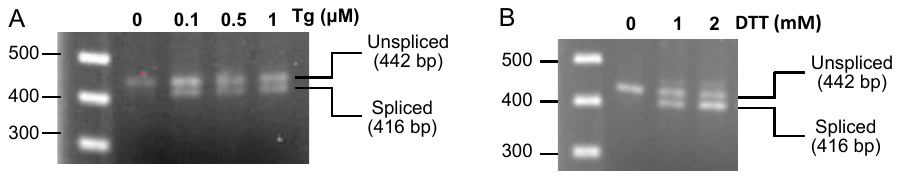

**Figure S4.** **Thapsigargin and DTT induce *XBP1* splicing in K-562 cells.** K-562 cells were treated with the indicated concentrations of Tg for 3 h (A) or DTT for 2 h (B) and *XBP1* splicing was determined by semi-quantitative PCR.

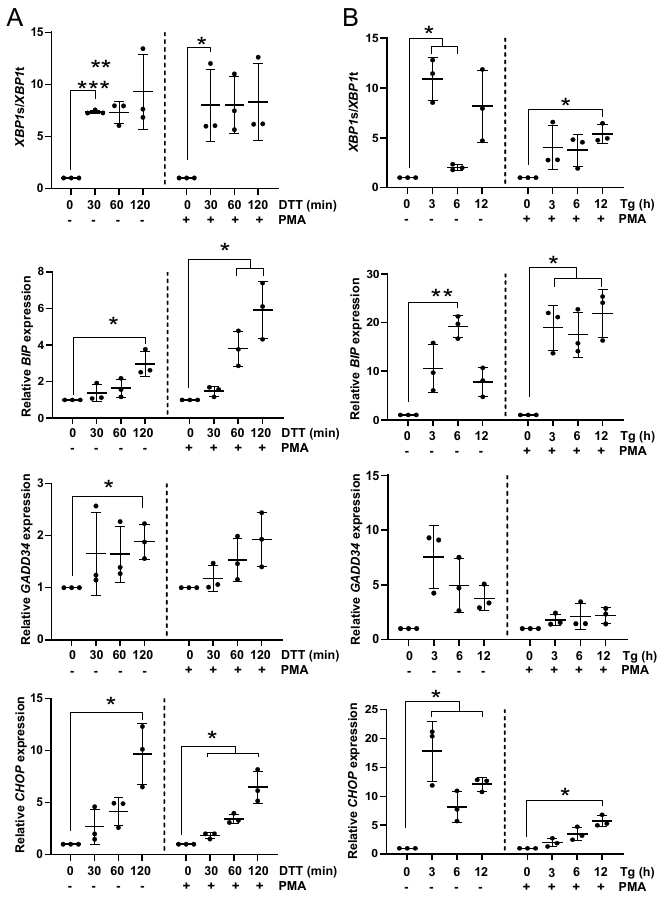

**Figure S5.** **UPR activation in immature and mature MEG-01 cells in response to DTT or thapsigargin (Tg).** Immature (no PMA) or mature (3 days after PMA treatment) MEG-01 cells were treated with 10 mM DTT for 0-2 h (A) or 0.1 µM Tg for 0-12 h (B). Expression of *XBP1*s and *XBP1*t, *BIP, GADD34* and *CHOP*, mRNA was quantified by RT-qPCR. Expression is relative to 0 h. N=3 ± SD. **P* < 0.05, ***P* < 0.01 (one sample t-test).

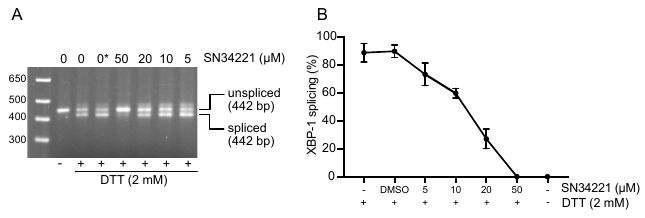

**Figure S6. SN34221 inhibits IRE1α endonuclease activity.** K-562 cells were pretreated with 0-50 μM SN34221 for 30 min, followed by 2 mM DTT for 6 h to induce the UPR. **A.** cDNA samples amplified using *XBP1* primers and analyzed by agarose gel electrophoresis. The product sizes of un-spliced and spliced *XBP1* are shown. *DMSO. **B.** % *XBP1* splicing determined from densitometry of agarose gels. N = 2 ± SD.

**Table S1. Caspase activation during PMA-induced maturation of K-562 cells**

| **Treatment** | **Relative caspase 3 activity** |
| --- | --- |
| untreated | 1 |
| 0.5 nM PMA, 1 day | 0.6 |
| 0.5 nM PMA, 2 days | 1.0 |
| 0.5 nM PMA, 3 days | 1.2 |
| 10 mM DTT, 22 h | 5.0 |

**Table S2. RT-qPCR primers used in the study.**

| **Gene** | **Forward primer (5’-3’)** | **Reverse primer (5’-3’)** | **Product size (bp)** | **Species** |
| --- | --- | --- | --- | --- |
| *CHOP (DDIT3)* | GGAGCATCAGTCCCCCACTT | TGTGGGATTGAGGGTCACATC | 101 | Human |
| *BIP (GRP78)* | TGACATTGAAGACTTCAAAGCT | CTGCTGTATCCTCTTCACCAGT | 116 | Human |
| *GADD34 (PPP1R15A)* | CCCAGAAACCCCTACTCATGATC | GCCCAGACAGCCAGGAAAT | 102 | Human |
| *ERdj4 (DNAJB9F)* | GATACACTTGGACACAGTGC | CTACTGTCCTGAACAGTCAG | 414 | Human |
| *HERPUD1* | AACGGCATGTTTTGCATCTGGTGTG | CAGGGGAAGAAAGGTTCCGAAG | 180 | Human |
| *XBP1*t | GGCATCCTGGCTTGCCTCCA | GCCCCCTCAGCAGGTGTTCC | 75 | Human |
| *XBP1*s | CGCTTGGGGATGGATGCCCTG | CCTGCACCTGCTGCGGACT | 101 | Human |
| *GAPDH* | GCTCTCTGCTCCTCCTGTT | CATGGTGTCTGAGCGATGTG | 79 | Human |
| *HPRT1* | CCTGGCGTCGTGATTAGT | ACCCTTTCCAAATCCTCAGC | 89 | Human |
| *GATA1* | CACGACACTGTGGCGGAGAAAT | TTCCAGATGCCTTGCGGTTTCG | 140 | Human |
